# Unique RNA replication characteristics and nucleocapsid protein expression may explain differences in the replication capacity of SARS-COV-2 lineages

**DOI:** 10.1101/2024.05.14.594070

**Authors:** Isadora Alonso Corrêa, Marcos Romário Matos de Souza, Gustavo Peixoto Duarte da Silva, Anna Beatriz Sampaio Vianna Macedo Pimentel, Pedro Telles Calil, Marcela Sabino Cunha, Diana Mariani, Rodrigo de Moares Brindeiro, Sara Mesquita Costa, Maria Clara da Costa Simas, Victor Akira Ota, Elisa Cavalcante Pereira, Marilda Mendonça Siqueira, Paola Cristina Resende, Rafael Mello Galliez, Debora Souza Faffe, Rosane Silva, Terezinha Marta Pereira Pinto Castiñeiras, Amilcar Tanuri, Luciana Jesus da Costa

## Abstract

COVID-19 pandemic in Brazil was characterized by the sequential circulation of the SARS-CoV-2 lineages B.1.1.33, and variants Zeta (P.2), Gamma (P.1/P.1.*), Delta (B.1.617.2/AY.*), and Omicron (BA.*). Our research aimed to compare the biological traits of these lineages and variants by analyzing aspects of viral replication including binding, entry, RNA replication, and viral protein production. We demonstrated that the replication capacity of these variants varies depending on the cell type, with Omicron BA.1 exhibiting the lowest replication in the human pulmonary cells. Additionally, the nucleocapsid proteoforms generated during infection exhibit distinct patterns across variants. Our findings suggest that factors beyond the initial stages of virus entry influence the efficiency of viral replication among different SARS-CoV-2 variants. Thus, our study underscores the significance of RNA replication and the role of nucleocapsid proteins in shaping the replicative characteristics of SARS-CoV-2 variants.

**Author summary:** The COVID-19 pandemic was characterized by the emergence of different viral variants that presents specific properties such as response to antibodies, pathogenicity and detection by diagnostic tests. The circulation of these variants presented a particular pattern depending on the global geographic regions. Despite the cessation of the pandemic, as officially declared by the World Health Organization in 2023, new viral variants continue to emerge while aspects of the virus-cell interaction that contribute to the replication of these variants have not yet been completely understood. In our study, we compared the biological characteristics of SARS-CoV-2 variants that circulated in Brazil during the pandemic, verifying aspects of entry, viral replication and production of viral RNA and proteins. Our results indicate that Omicron BA.1 variant has reduced replication and protein production in human lung cells. We also observed that the viral nucleocapsid protein presents proteoforms that vary according to the variant. These differences could help to explain the differences observed in viral replication in human pulmonary cells.

## Introduction

The Severe Acute Respiratory Syndrome Coronavirus 2 (SARS-CoV-2) was identified in December 2019 and responsible for the coronavirus disease 2019 (COVID-19) pandemic. Despite its lower mutation rates compared to other RNA viruses, some factors contribute to the accumulation/selection of mutations within the viral genome such as: the widespread virus circulation, independent introduction events, host adaptive immune pressure, persistent infections, and the high immunization level^1,2,3^. The first major mutation that altered viral dynamics was the Spike (S) glycoprotein substitution D614G, which arose in March 2020 and was rapidly overcoming the original Wuhan virus^4^. Studies demonstrated that G614 viruses replicate more than D614 in the superior airways and, in hamster models and in competition assays, could overcome D614 viruses even when these were the major population^5,6^. The diversity of SARS-CoV-2 genomes led to the WHO’s classification as variants of interest (VOIs) and variants of concern (VOCs), which possess genetic changes that could affect viral transmissibility, disease severity, and immune escape^7^.

The VOCs Alpha (B.1.1.7) and Beta (B.1.351) emerged in the United Kingdom and South Africa, respectively, in 2020. The VOC Delta (B.1.617.2/AY.*), first described in October 2020 in India, quickly spreading around the world and accumulated more mutations in the S gene than the previous variants^8^. However, in November 2021, Delta was replaced by a new VOC that emerged in Botswana and South Africa named Omicron (BA.1). In January 2022, Omicron BA.1 dominated the pandemic scenario even in countries with high vaccination rates^9,10^. This variant presented over 30 substitutions in the S protein^11^ and currently, continues to evolve^9^.

After the first case confirmed in Brazil, in late February 2020, multiple viral introductions were documented in the country with the predominance of two major lineages, B.1.1.28 and B.1.1.33^12^. From this lineages derived the VOC Gamma (P.1/P.1.*) in December 2020 and, by January 2021, became the most prevalent variant in the country and was associated with a high mortality rate^13^, high viral loads and transmissibility^14,15^. VOI Zeta (P.2), also derived from B.1.1.28 and B.1.1.33, emerged around October 2020 and was associated with an increase in infection rates in the country^16,17^.

SARS-CoV-2 circulation is highly dynamic, and the mechanisms driving the spread and infectiousness of VOCs and VOIs still need to be clarified. For VOCs Delta and Gamma, lower cycle thresholds (Ct values) were reported in patients positive for SARS-CoV-2^18,19^, and Delta was associated with increased disease severity^20^. Despite its high transmissibility, several studies indicate that VOC Omicron BA.1 has impaired replication in pulmonary cells^21,22^, resulting in lower pathogenicity in mouse and hamster models than previous VOCs^23,24^. Also, nasal samples from Omicron-infected patients showed lower viral loads when compared to Delta infections^25,26^. However, Omicron presents faster replication rates in nasal epithelium and human bronchi^22,24^.

Hence, improving the knowledge about the biological characteristics and replication dynamics of SARS-CoV-2 variants remains crucial to understanding the potential of this pandemic virus to adapt to new scenarios and guide better strategies to prevent the emergence of new VOCs. The present study compares the biological characteristics of SARS-CoV-2 variants B.1.1.33, Zeta, Gamma, Delta, and Omicron BA.1. Our work indicates that Omicron BA.1 has the lowest replication compared to previous variants in a human pulmonary cell line. Besides, our results highlight diverse aspects of viral replication for each variant, including binding, entry, RNA replication, and viral protein production. Collectively, we demonstrate the importance of the RNA replication step and the role of Nucleoprotein (N) in shaping the replicative characteristics of the SARS-CoV-2 variants.

## Results

### Distinct waves of SARS-CoV-2 infection in Brazil

Distinct waves of infection occurred in Brazil since February 2020, marking the emergence, circulation, and extinction of SARS-CoV-2 VOI and VOC. Analysis of sequencing data obtained from Brazilian samples and compiled by Fiocruz Genomic Surveillance Network (23) demonstrated that lineages B.1.1.28 and B.1.1.33 were present at the pandemic’s beginning and co-circulated predominantly up to September 2020. At that time, the Zeta variant emerged and prevailed being replaced by VOC Gamma. Gamma rapidly dominated the Brazilian scenario, causing a second wave of infections that began in March 2021 (Figure 1A). After Gamma’s ascendence, the VOC Delta was introduced in July 2021. Delta remained prevalent from October to December 2021 despite a decrease in confirmed cases. However, the introduction of Omicron (BA.1) caused a fourth wave of infections, leading to the highest number of reported cases until June 2022 (Figure 1A).

**Figure 1:**
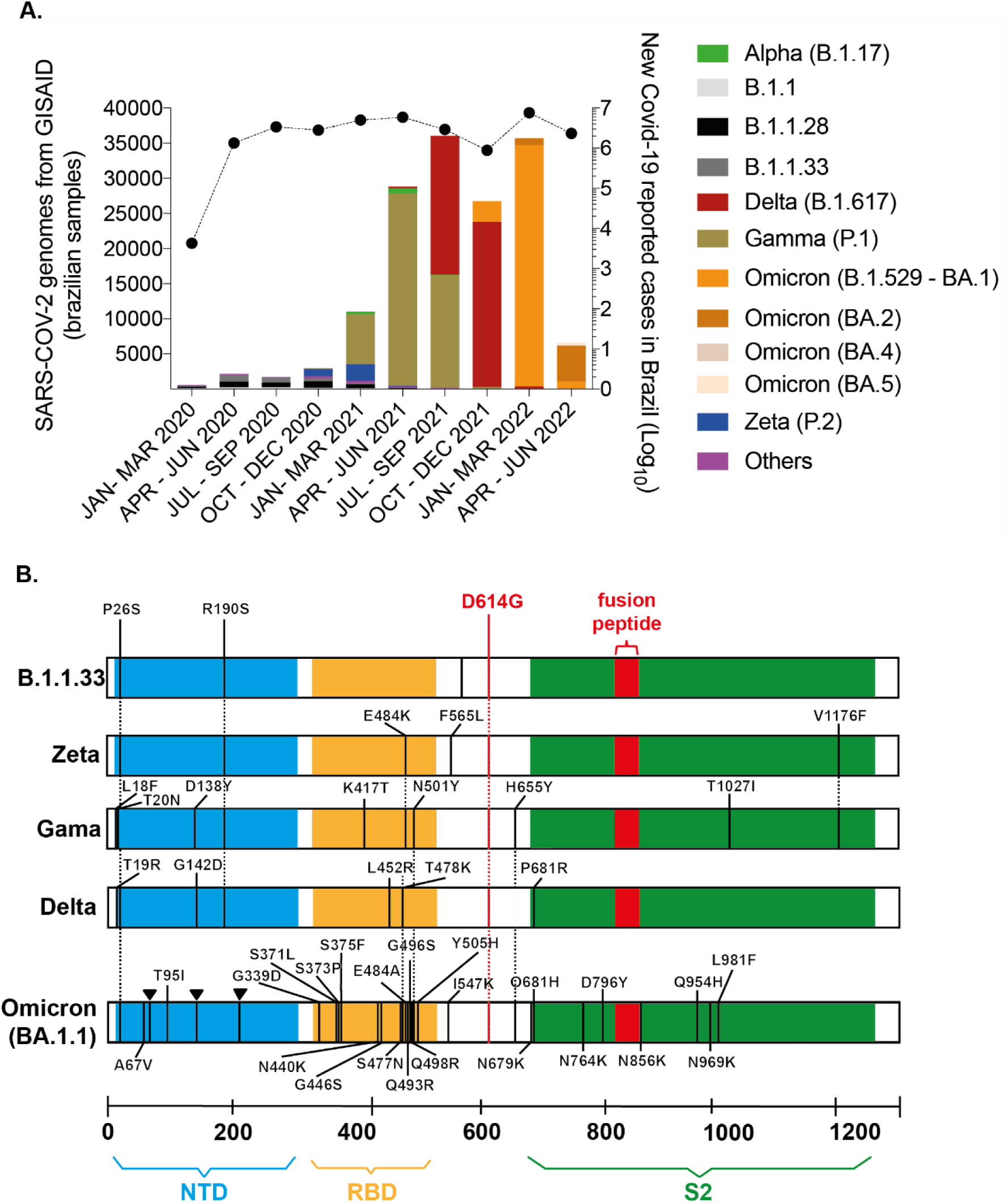
SARS-CoV-2 variants were associated with distinct waves of infections in Brazil. **A-** Sequences derived from NGS sequencing of SARS-CoV-2 Brazilian samples were compiled by Fiocruz Genomic Network from January 2020 to June 2022. The left y-axis indicates the absolute number of deposited sequences. Each color represents a different viral lineage. The right y-axis shows the total number of Covid-19 cases in Brazil registered by the Covid global data from WHO during the same period (black line). **B-** Spike (S) region representation for B.1.1.33, Zeta, Gamma, Delta, and Omicron (BA.1) genomes obtained by NGS sequencing of viral stocks generated after SARS-CoV-2 isolation. The defining mutations of each lineage are indicated according to protein domains: N-terminal domain (NTD) in blue; receptor binding domain (RBD) in yellow; S2 subunit in green; and the fusion peptide in red.

Our group continuously isolated and characterized the main circulating SARS-CoV-2 lineages from samples of patients attending NEEDIER. B.1.1.33, Zeta, Gamma, and Delta viruses were successfully isolated in VeroE6 cells. NGS sequencing was performed for these variants along with BA.1, and characteristic polymorphisms at the S gene and other genomic regions confirmed viral lineages (Figure 1B and Supplementary Fig. 1). Viral stocks were generated from passages 2 to 4, depending on the variant, and few mutations were selected during cell culture passages in VOCs Gamma (Orf1a and ORF3a), Delta (Orf1a), and BA.1 (Orf1a and Spike RBD domain) (Supplementary Fig.1).

### The advantages in the binding step did not correlate with the entry capacity of SARS-CoV-2 variants

Although all variants efficiently bound to the surface of Vero E6 and Vero/hACE-2/hTMPRSS2 cells, B.1.1.33 presented the highest binding capacity when compared to Gamma, Delta and BA.1 (Figures 2A and 2B - top) in Vero/hACE-2/hTMPRSS2 cells. When compared to all other variants, BA.1 had the lowest ability to bind to Vero/hACE-2/hTMPRSS2 cells (Figure 2B) while was the most efficient to bind to the Calu-3 cell surface (Figure 2C).

**Figure 2:**
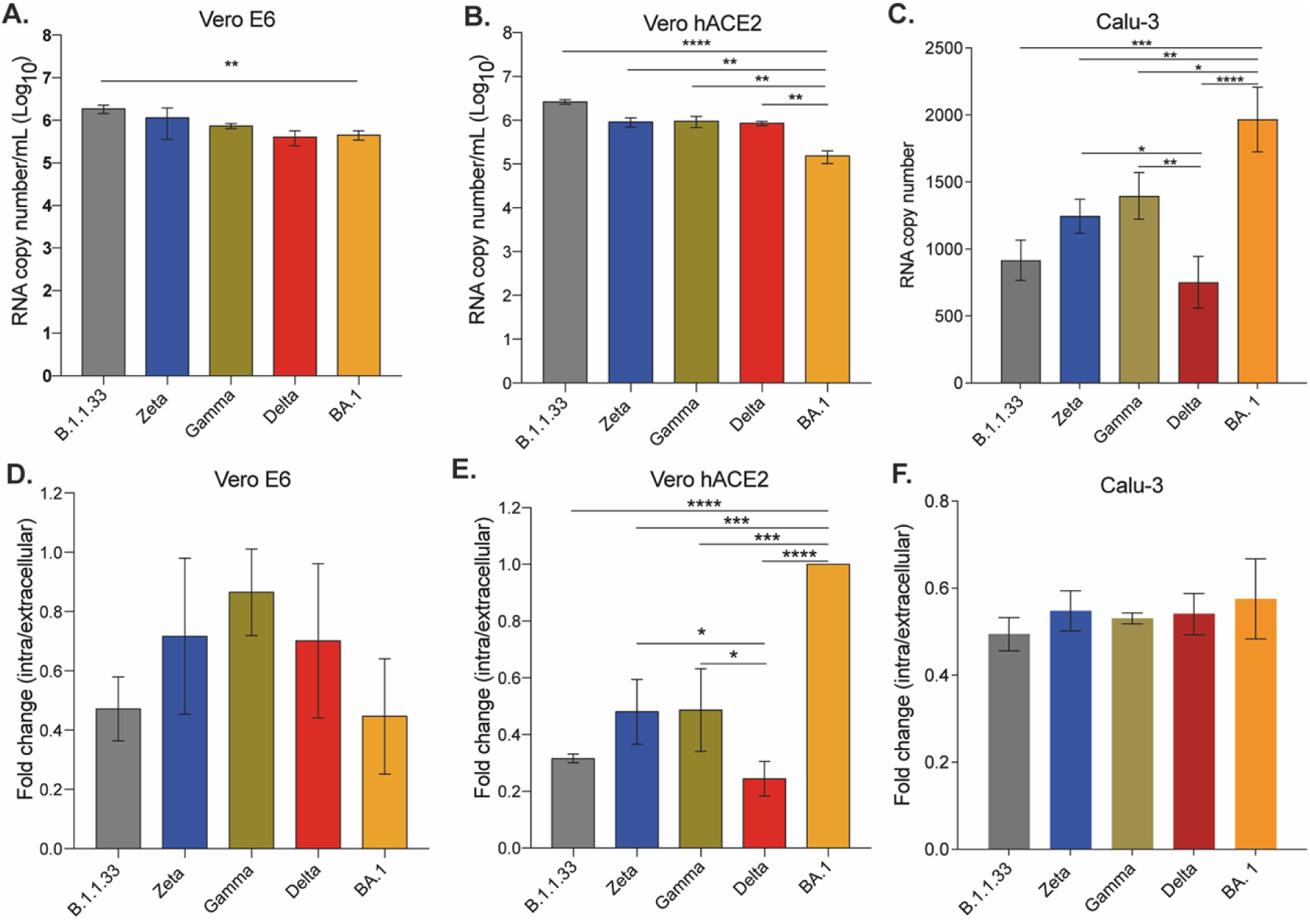
SARS-CoV-2 variants presented differential binding and entry capacities. Vero E6 **(A)**, Vero/hACE-2/hTMPRSS2 **(B)**, and Calu-3 **(C)** cells were incubated with the same copy number of RNA of each viral variant for 1 hour at 4°C to allow viral binding. After adsorption, cells were harvested for RNA extraction and SARS-CoV-2 detection and quantification by RT-qPCR. For entry assay, Vero E6 **(D)**, Vero/hACE-2/hTMPRSS2 **(E)**, and Calu-3 **(F)** cells were incubated for 1 hour at 4°C and after adsorption, the inoculum was replaced by DMEM with 10% FBS and cells were incubated at 37°C with 5% CO2 for one more hour to allow entry. The efficiency of viral entry was calculated by the ΔCt between cellular lysates obtained after incubation at 37°C and at 4°C. Data are shown as mean±s.d. n=3. Statistical analysis was performed using one-way analysis of variance (ANOVA) followed by Dunnettés multiple comparisons test; p<0.05 was considered statistically significant. *p<0.05; **p<0.01; ***p<0;001; ****p<0.0001.

However, the greater binding ability observed for some variants in certain cell lines did not correlate to their entry capacity. For Vero E6 cells, despite the greater biding efficiency of B.1.1.33, there was no difference in entry among all viral variants analyzed (Figure 2D). On the other hand, despite the lower binding capacity of BA.1 on Vero/hACE-2/hTMPRSS2 cells, this variant presented the highest entry capacity compared to all other variants analyzed (Figure 2E). In this cell line, the Delta variant presented significantly lower entry capacity compared to Zeta, Gamma, and BA.1 (Figure 2E). Again, the highest ability of BA.1 to bind to Calu-3 celĺs surface did not correlate with a better entry capacity, no difference in entry ability was observed amongst the variants in Calu-3 cells (Figure 2F). The expression of ACE-2 on Vero/hACE-2/hTMPRSS2 cells was similar to Calu-3 (Supplementary Fig. 2). According to our results, the SARS-CoV-2 efficiency of binding and entry did not depend on levels of ACE-2 expression or the presence of TMPRSS2. In general, the native SARS-CoV-2 variants bound with the same efficiency to the surface of these cell lines, a difference in entry was only observed in the cell line that heterologously expressed TMPRSS2 and ACE-2.

### SARS-CoV-2 BA.1 had the highest replication capacity both in Vero-hACE and Vero E6 cells

Vero/hACE-2/hTMPRSS2 cells were infected with the SARS-CoV-2 variants, and total and infectious viral production was measured over time. BA.1 and Gamma had the highest titers of infectious viral progeny production up to 48 hpi (Figure 3A). Overall, BA.1 had the highest production of viral infectious progeny (Figure 3B). The same pattern was observed for the total viral progeny production when the genomic RNA was quantified (Figure 3B). Interestingly, all variants had similar amounts of infectious virus and total viral particle production at 72 hpi (Figures 3A and 3B), suggesting that B.1.1.33 and Delta had a delayed production curve. The particle-to-PFU ratio indicates that, except for B.1.1.33 and Zeta, which had an overall higher ratio, all VOCs remained similar, suggesting that SARS-CoV-2 variants before VOCs led to a higher production of defective viral particles (Figure 3C).

**Figure 3:**
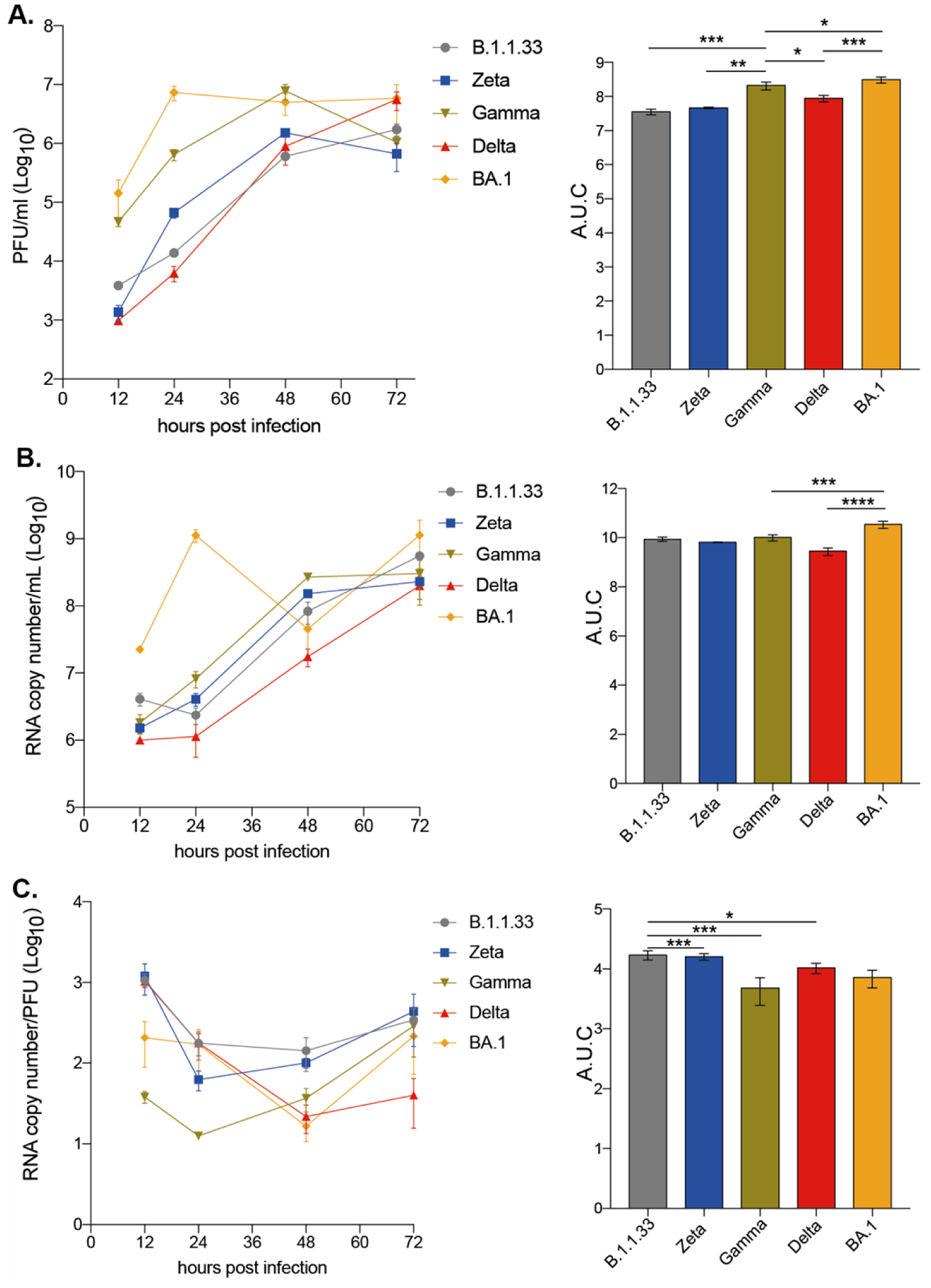
BA.1 variant showed greater infection capacity in Vero-hAce2/h-TMPRST2 cells. Cells were infected with viral variants at MOI 0.1 and incubated at 37°C with 5% CO2 for 12, 24, 48, or 72 hours to evaluate viral replication kinetics. **A-** Infectious viral progeny quantification by plaque assay at indicated time points. **B-** Viral RNA copy number quantification by RT-qPCR at indicated time points. **C-** Particle to plaque-forming unit ratio (P:PFU) obtained as a measurement of viral infectivity. Viral titers **(A)**, RNA copy numbers **(B)**, and P:PFU **(C)** are also depicted as areas under the curve (AUC). Data are shown as mean±s.d. n=3. Statistical analysis of AUC was performed using one-way ANOVA followed by Dunnettés multiple comparisons test; p<0.05 was considered statistically significant. *p<0.05; **p<0.01; ***p<0;001; ****p<0.0001.

We observed that BA.1 followed by Gamma had the highest total and infectious viral progeny production in Vero E6 up to 48 hpi (supplementary figure 3A). However, BA.1 titers peaked at 48 hpi and maintained at 72 hpi, while Delta and B.1.1.33 peaked at 72 hpi., confirming its delayed production curve. Overall, Zeta was the least replicating variant in Vero E6 cells (Supplementary Figures 3A and 3B), correlating with an increasing particle-to-PFU ratio (Supplementary Figure 3C). Gamma and Delta, which achieved high viral infectious titers in 48 and 72 hpi, respectively, had the lowest particle-to-PFU ratio (Supplementary Figure 3C).

The results from infection of Vero E6 cells and Vero cells expressing human ACE-2 and TMPRSS2 indicate the greatest replication capacity for BA.1, followed by Gamma. Moreover, Delta replicated more efficiently than B.1.1.33 and Zeta variants, suggesting that in Vero cells, the more recent variants have a replication advantage, which solely in the case of BA.1 in Vero-hACE cells could be explained by its greater efficiency for cell entry. The same behavior observed in Vero E6 cells points out some advantages for these VOCs beyond the cell attachment/entry replication steps.

### BA.1 was less infectious in a human pulmonary cell line compared to previous SARS-CoV-2 variants

The infection of Calu-3 cells with the SARS-CoV-2 variants showed that BA.1 had the lowest replication capacity up to 48 hpi. Levels of infectious viral progeny and total viral production were 1,000- to 15-fold and 10-fold lower, respectively, for BA.1 when compared to all other variants tested between 24-48 hpi (Figures 4A and 4B). B.1.1.33, Zeta, Gamma, and Delta presented similar infectious abilities in this cell line. AUC analysis confirmed that BA.1 had the lowest infectious virus progeny (Figure 4A). Levels of sgRNA, as a surrogate for RNA replication, confirmed that BA.1 replicated less during the entire course of infection (Figure 4C). However, it could reach similar levels of infectious virus production to the other variants at 72 and 96 hpi, when those already show decreasing infectious titers (Figure 4A). Gamma and Delta had significantly higher levels of sgRNA compared to BA.1 from 6 to 24 hpi (Figure 4C). As for Zeta and Gamma, no differences were observed when compared to B.1.1.33 (Figure 4D). Overall, Gamma and Delta, the highest replicating variants in this cell line, had the highest particle-to-PFU ratio (Figure 4E). The highest replication rates of B.1.1.33, Zeta, Gamma, and Delta were associated with a significant percentage of cell-death induction from 48 hpi, especially for the Delta variant (Figure 4A and 4F). In contrast, for BA.1, more than 50% of cell survival was maintained up to 96 hpi., indicating that cell-induced death correlates with the extension of SARS-CoV-2 replication (Figure 4F). Altogether, these results demonstrated that the BA.1 variant does not replicate efficiently in a human pulmonary cell line compared to previous SARS-CoV-2 variants.

**Figure 4:**
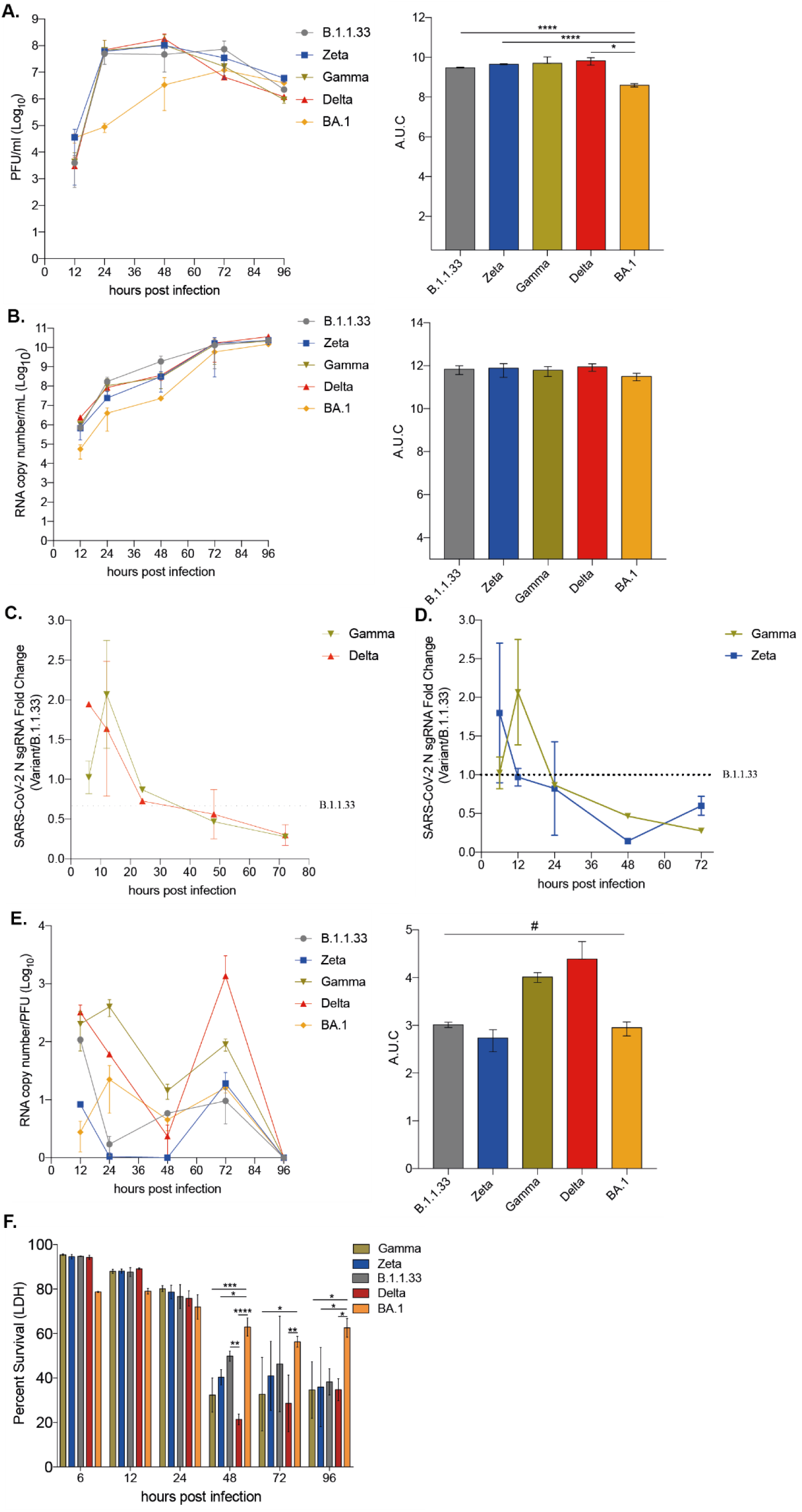
Replication of SARS-CoV-2 BA.1 variant was reduced compared to previous variants in pulmonary cells. Calu-3 cells were infected with viral variants at MOI 0.1 and incubated at 37°C with 5% CO_2_ for 12, 24, 48, 72, or 96 hours to evaluate viral replication kinetics. **A-** Infectious viral progeny quantification by plaque assay at indicated time points. **B-** Viral RNA copy number quantification by RT-qPCR at indicated time points. **C-** Nucleocapsid subgenomic RNA (sgRNA) quantification by RT-qPCR of variants Gamma, Delta, and BA.1 represented as fold change in relation to B.1.33 variant. **D-** Nucleocapsid subgenomic RNA (sgRNA) quantification by RT-qPCR of variants Zeta and Gamma represented as fold change in relation to B.1.1.33 variant. **E-** Particle to plaque-forming unit (P:PFU) was obtained as a measurement of viral infectivity. **F-** Percentage of cell viability measured by LDH released in the cell culture supernatant at the indicated time points after infection. Vitral titres (**A**), RNA copy numbers (**B**), and P:PFU (**E**) are also depicted as area under the curve (AUC). Data are shown as mean±s.d. *n*=3. Statistical analysis of AUC was performed by one-way anova followed by Dunnettés multiple comparisons test. Statistical analysis of sgRNA quantification was performed by unpaired *t* tests. *p*<0.05 was considered statistically significant. *p<0.05; **p<0.01; ***p<0;001; ****p<0.0001; # no statistically significant difference was observed when analyzed by one-way anova.

A study by Hui and collaborators^29^ demonstrated that BA.1 presents different replicative kinetics according to the temperature of incubation and the cells from the upper and lower respiratory tract. When Calu-3-infected cells were maintained at 35°C we still observed a lower replication rate for BA.1 compared to the other variants after 24 hours of infection albeit to a lower extent at 35°C. However, the BA.1 infectious titer had a 0.5 log10 increase at 35°C when compared to 37°C, while the other variants present a statistically 2-3 log10 decrease in viral titers when replicating at 35°C (supplementary figure 4). These results indicate that lowering the temperature alleviated the disadvantage of BA.1 replication in Calu-3 cells.

### Direct competition assay confirmed the replicative disadvantage capacity of the SARS-CoV-2 Omicron variant in Calu-3 cells but not in Vero/hACE-2/hTMPRSS2

Competition assays using two SARS-CoV-2 variants to co-infect Calu-3 and Vero/hACE-2/hTMPRSS2 cells in different proportions were performed, followed by an NGS or RT-qPCR assay to detect the viral populations produced. In Calu-3 cells infected with BA.1 and Gamma, the population analysis after NGS indicated that BA.1 is the major population only when used nine times more than Gamma (9:1 proportion) at 24 hpi (Figures 5A-C). The Gamma variant suppressed BA.1 in all the remaining conditions. Further, Delta outgrew BA.1 even with a 9-fold higher BA.1 inoculum. In all BA.1 to Delta proportions, the frequency of the BA.1 genotype in the viral progeny population was around 10% (Figures 5D–F).

**Figure 5:**
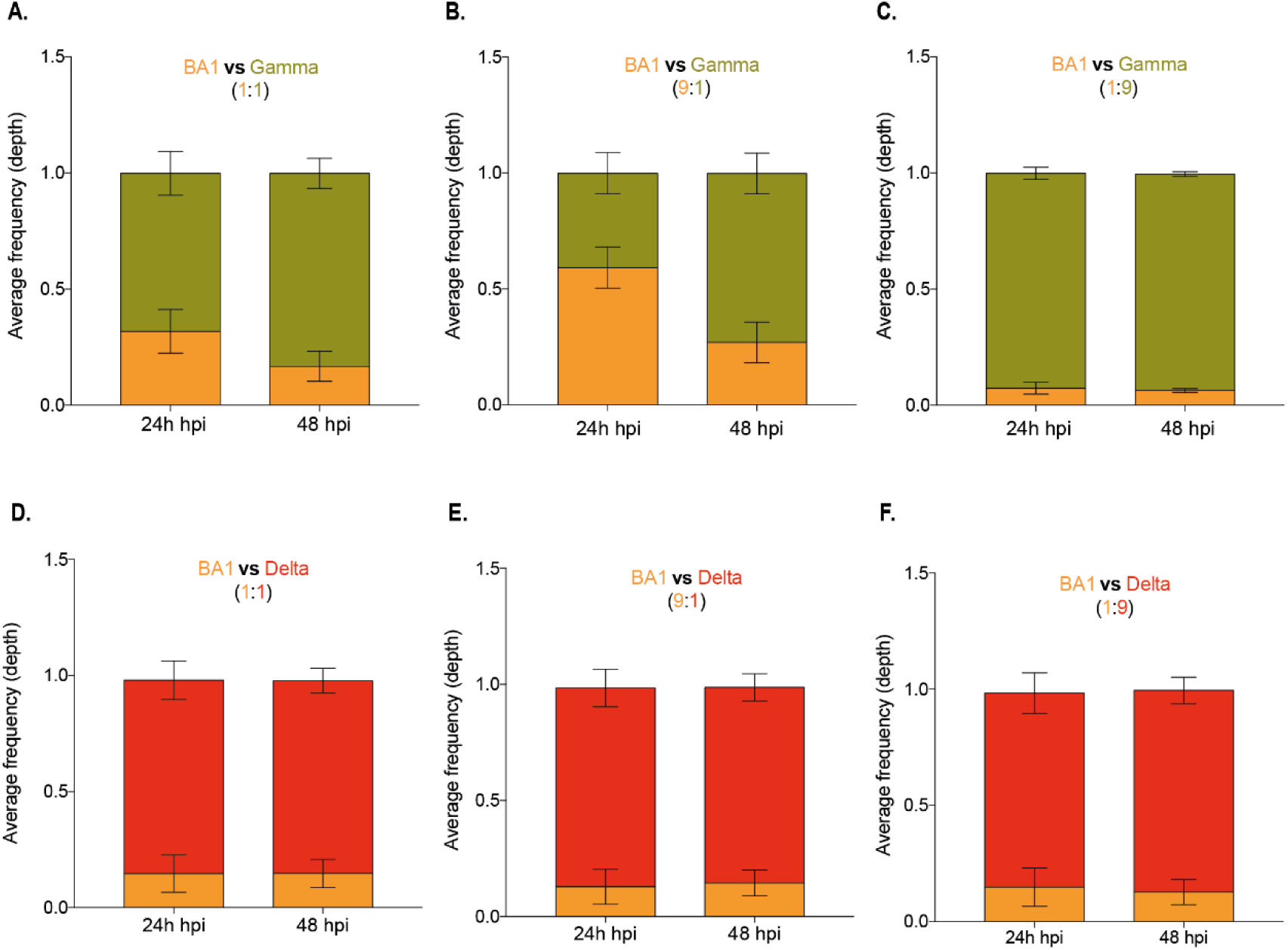
Competition assay in Calu-3 cells confirmed the lower replication of SARS-CoV-2 BA.1 variant. Calu-3 cells were co-infected with three different proportions of SARS-CoV-2 variants, resulting in a final MOI of 0.1 PFU. The infection was performed with variants Omicron BA.1 and Gamma at 1:1 **(A)**, 1:9 **(B)**, and 9:1 **(C)** proportions and with variants Omicron BA.1 and Delta at 1:1 **(D)**, 1:9 **(E)**, and 9:1 **(F)** proportions. Total RNA was extracted from culture supernatant 24- or 48-hours post-infection and subjected to NGS sequencing. The depth of each genomic region from viral lineages was plotted as the average frequency of detected sequences. Bar colors represent viral lineages BA.1 (yellow), Gamma (green) and Delta (red). Data are shown as mean±s.d. *n*=3.

Co-infection in Vero/hACE-2/hTMPRSS2 cells demonstrated that Gamma only dominated over BA.1 at 24 hpi at a 1:1 proportion and at both 24 and 48 hpi at a 9-fold excess of the Gamma inoculum (Supplementary Figures 5A–C). While for Delta and BA.1 co-infection, BA.1 only dominated at a 9-fold excess of the BA.1 inoculum (Supplementary Figures 5D–F). These data confirmed the previous replication kinetics results in Vero/hACE-2/hTMPRSS2 cells, where although the BA.1 infection led to a higher infectious virus progeny production, the overall difference for Gamma and Delta was 0.5 – 1.0 log10 (Figure 3A). In contrast, in Calu-3 cells, BA.1 replication was markedly reduced compared to Gamma and Delta (2 log10 overall difference – Figure 4A).

The same assay was performed with Zeta, Gamma, and Delta variants infecting Calu-3 cells and revealed by RT-qPCR. In direct competition for Gamma and Delta, the latter could not suppress Gamma replication at any proportion (supplementary figure 6A – C). At 24 hpi, even at the 9:1 Delta to Gamma proportion, only 30% of genome detection was from Delta (supplementary figure 6B). As for the Zeta and Gamma competition (supplementary Figures 6D–F), the Zeta variant had a replicative advantage only at the 9:1 Zeta to Gamma proportion at 24 hpi. Both variants were equally detected at 48 hpi (supplementary figure 6E). Gamma outgrew Zeta in all the other conditions (supplementary Figures 6D and 5F), demonstrating the highest fitness of Gamma compared to the previous circulating variant Zeta and the replacing Delta.

Altogether, these results indicate that viruses that became predominant in the pandemic scenario and replaced variants that circulated previously do not necessarily present replicative advantages in cell culture. This suggests that other factors besides virus fitness are related to replacing a circulation SARS-CoV-2 variant with another dominant variant during the pandemic.

### The size and number of RNA replication sites in infected cells varies amongst variants early in the SARS-CoV-2 replication cycle

To investigate possible factors involved in the difference in VOC’s fitness in tissue culture, we analyzed the early and late viral RNA replication during the viral replicative cycle in Calu-3 cells using the dsRNA as a marker. dsRNA was visualized as a predominant perinuclear puncta distribution early at infection (8 hpi) (Figure 6A). The average puncta per infected cell ranged from 25–250, with the lowest amount observed in Delta- and BA.1-infected cells and the highest in Zeta-infected cells (Figure 6B). All variants’ average puncta area size was similar, varying from 75-90μm^2^, except for the Delta, which had a puncta area approximately 2-fold higher (Figure 6C). Later during the SARS-CoV-2 replication cycle the dsRNA puncta were readily visualized and more distributed through the infected cell’s cytoplasm except for BA.1, that still maintained a predominant perinuclear distribution (Figure 6D). At this time point, both the average of puncta per infected cell (Figure 6E) and the average puncta area size (Figure 6F) increased, except for Gamma, which had a marked reduction in the average of puncta per infected cell (Figure 6E). No difference was observed for the average puncta area size of Delta and the other variants at this time point (Figure 6F). These results demonstrated that a higher number of RNA replication sites and the size of these sites formed early in infected cells could contribute to a higher replication rate for Zeta and Gamma and for Delta, respectively. Indeed, Zeta and Delta had the highest percentage of cells actively replicating SARS-CoV-2 at 8 hpi. The increased size of the RNA replication organelle for Delta resulted in a higher rate of cells maintaining RNA replication up to 24 hpi (Supplementary Figure 7).

**Figure 6:**
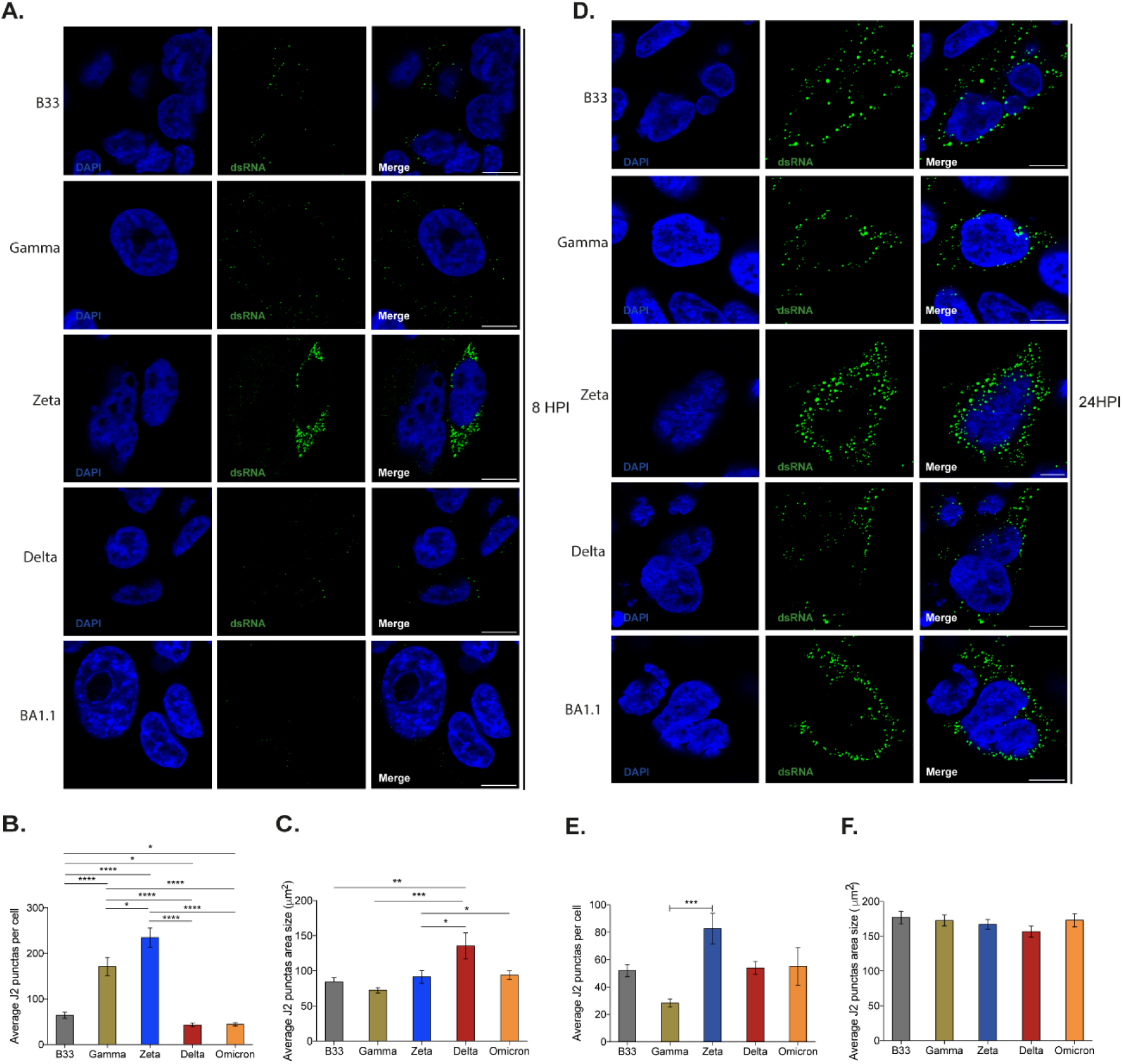
Early and late SARS-CoV-2 VOC’s replication in Calu-3 cells analysis through dsRNA immunofluorescence staining. **A and D** - Representative images of confocal microscopy analysis of VOC’s dsRNA accumulation at 8 and 24 hours post-infection, respectively. **B and E** - Quantification of dsRNA punctas (average) per cell at 8 and 24 hours post-infection, respectively. **C and F** - Quantification of dsRNA punctas area size (in µm^2^) at 8 and 24 hours post-infection, respectively. Data are shown as mean±s.d. *n*=30. Scale bar = 10 µm. Statistical analysis was performed by unpaired *t* tests. *p<0.05, **p<0.01, ***p<0.001, ****p<0.0001 by unpaired t-test.

### S and Nucleocapsid (N) viral proteins accumulated to higher levels in VOC Delta Calu-3-infected cells at 24 hpi

We analyzed the expression levels of the main SARS-CoV-2 structural proteins, S and N, in infected Calu-3 cells at 24 and 48 hpi. The Delta variant presented the highest full-length (FL) and total (FL + processed S1) S glycoprotein levels at 24 hpi, followed by Zeta. However, at 48 hpi, Delta had a reduction in FL and total S and the B.1.1.33 becomes the variant with higher protein production. At the same time point, BA.1 had an increase in FL S (Figures 7A–C).

**Figure 7:**
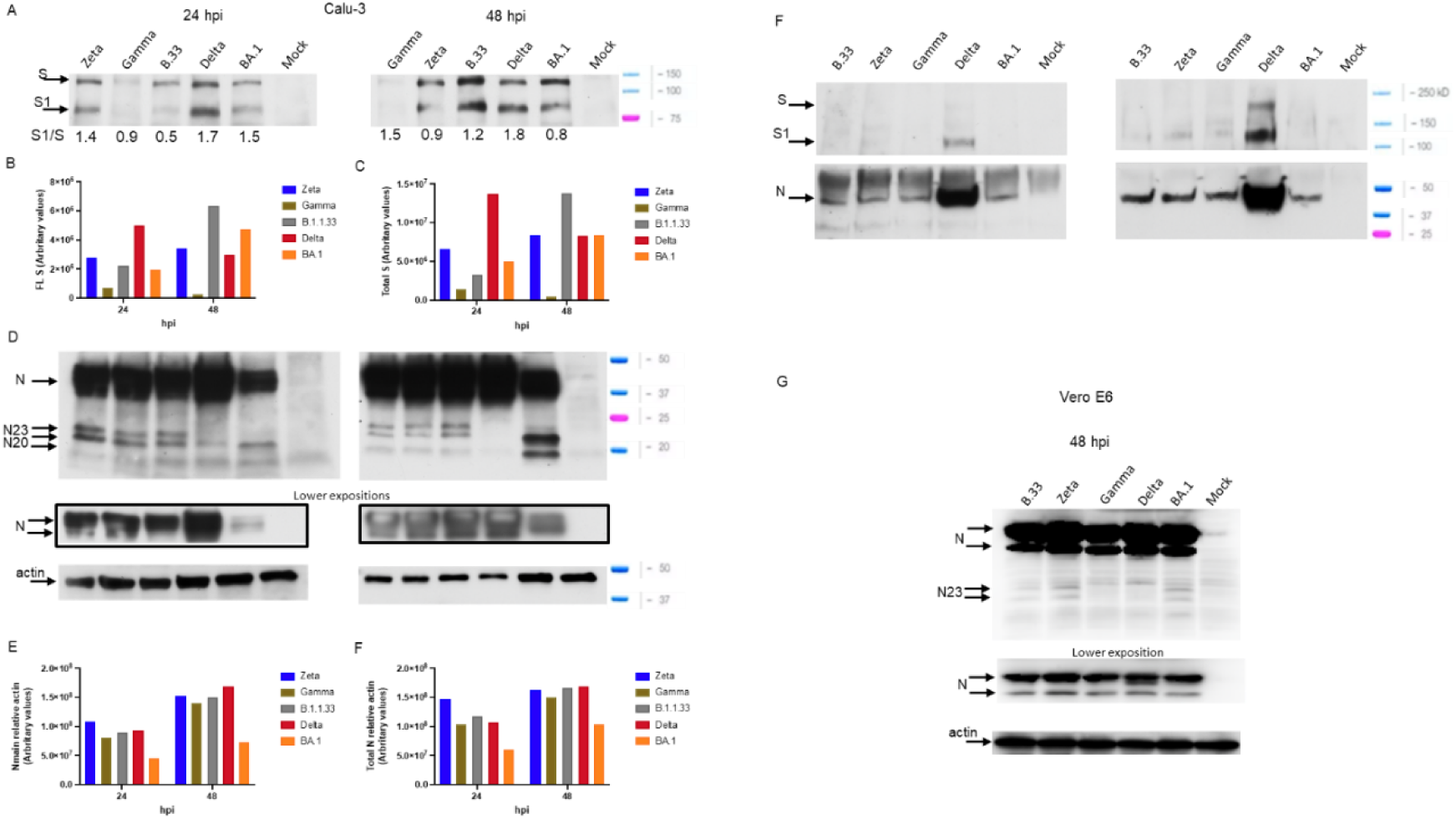
Differential levels of viral full-length and processed proteins accumulated in cells infected with SARS-CoV-2 variants. **(A)** - Calu-3 cells were lysed 24- or 48-hours post-infection and submitted to SDS-PAGE/WB with antibodies against viral S. Arrows indicated the full-length protein and its isoforms. Expressions of full-length (FL) S protein **(B)**, total S protein (full-length plus processed S1) **(C)**, N protein **(D)**, full-length N protein **(E)**, and total N protein (full-length plus isoforms) **(F)** were quantified by ImageJ and normalized by β-actin from arbitrary values. Arrows indicated the full-length protein and its isoforms. **(G)-** Cell-free supernatants submitted to SDS-PAGE/WB with antibodies against viral S and N proteins. **(H)-** Vero/hACE-2/hTMPRSS2 cells were infected with SARS-CoV-2 variants. After 48 hours of infection, cells were lysed and submitted to SDS-PAGE/WB with antibodies against viral N protein and human β-actin.

As previously demonstrated^30^, Delta had the highest S processing rate, as measured by the S1/S ratio, at both times. In contrast, BA.1 and Zeta S processing rates are reduced compared to Delta’s, while B.1.1.33 S processing rates increased at 48 hpi. (Figure 7A). Spike protein levels were low in Gamma-infected cells and decreased over time (Figures 7A–C), which did not correlate with the high levels of infectious progeny produced at both time points (Figure 4A). Spike protein detection for Gamma-infected cells did not improve with longer membrane exposition (Supplementary Figure 8).

The N protein levels were highest in Delta-infected cells at both time points and 2-fold lower in BA.1 infected cells than other variants (Figure 7D). Three extra specific smaller N bands were detected in infected cells for all variants except for the Delta in 48 hpi: two of 23 kDa and one of 20 kDa, which was more apparent at 48 hpi. (Figure 7D). For Zeta, Gamma, and B.1.1.33 altogether these alternative forms of N represented 4 -10% of the total N protein at both time points, however, for BA.1 it increased from 2 to 30% from 24 to 48 hpi. (Figure 7D -F), increasing the total N levels in BA.1 infected cells (Figure 7F). The highest levels of structural SARS-CoV-2 proteins in Delta infected cells reflected the highest levels of its proteins in the cell-free supernatants, with a 10- and 5-fold excess for S and N, respectively when compared to Zeta, Gamma, and B.1.1.33 and 12- and 7-fold excess for S and N, respectively, when compared to BA.1 (Figure 7G). For the S glycoprotein, we predominantly detected the processed S1 domain, while the extra N forms detected in the infected cell’s lysates were not observed in the cell-free supernatants.

These results demonstrate that the differences observed in infectious virus production in Calu-3 cells amongst these SARS-CoV-2 variants could be partially explained by the different levels of viral structural protein synthesis. Analyzing the N protein content in Vero-hACE2/hTMPRSS2 infected cells, in which these variants replicate at equivalent efficiencies (Figures 4A and 6), we did not observe substantial differences (Figure 7H). The same 23 kDa smaller N bands detected in Calu-3 infected cells were also detected in Vero-hACE2/hTMPRSS2 albeit in lower levels for BA.1 (Figure 7H).

## Discussion

This study characterized the replication capacity of SARS-CoV-2 variants that circulated during the COVID-19 pandemic, including an ancient D614G virus (B.1.1.33), one VOI (Zeta), and 3 VOCs (Gamma, Delta, and BA.1), that sequentially replaced each other in the population. We demonstrated that the replication capacity of these variants depended on the cell type, for instance, Zeta, Gamma, and Delta replicated better in a human lung epithelial cell line. BA.1 instead replicated to highest titers in macaque epithelial kidney cells, independently of the TMPRSS2 presence. It has been demonstrated that BA.1 preferentially uses the endocytic entry route and is less sensitive to TMPRSS2 inhibitors^29,31,32^. Indeed, in our virus entry assays with native viruses, we observed a clear advantage for BA.1 in Vero-hACE2/hTMPRSS2 cells, possibly reflecting its more efficient use of the endocytic machinery. Previous studies reported a rate-limiting intermediate in the HIV fusion process that can be arrested by decreasing temperature^33,34^. If the S protein binding to its receptor is not rate-limiting, lowering the temperature could favor endocytosis by fusion constant reduction^35^. Thus, in our assay, the binding temperature (4°C) could have affected the S fusion dynamics with the cell membrane, favoring endocytosis and yielding BA.1 an advantage over the previous variants more prone to cell-fusion entry.

In direct competition assays, BA.1 was fitter than Gamma to replicate in Vero-hACE2/hTMPRSS2 cells up to 48 hpi., although Gamma prevailed at 24 hpi, thus reflecting the single-infection kinetics in this cell line. These results could be explained by the higher efficiency of BA.1 entry in these cells. However, the same pattern of single-infection kinetics was observed in Vero E6 cells, in which BA.1 and Gamma surface membrane attachment and cell entry were equivalent. Thus, other viral characteristics besides binding and entry could explain this replicative difference. On the other hand, although total and infectious virus titers were significantly higher for BA.1 than for Delta in Vero-hACE2/hTMPRSS2 cells, Delta was fitter to replicate in this cell line as indicated by the direct competition assays. These data suggest that viral factors beyond the early steps of virus entry affect the replication efficiency of these variants. Indeed, it has been demonstrated that S properties as binding to ACE2 and fusogenicity do not explain differences in replication amongst SARS-CoV-2 VOCs and VOIs^30^. However, Meng^32^ suggested that Omicron entry is impaired in cell lines highly expressing TMPRSS2, probably explaining the lower replicative capacity of this variant in native virus replication assays in various cell lines. In contrast, our data showed no significant differences amongst all variants in cell attachment and entry in Calu-3 cells or advantage of BA.1 entry in Vero-hACE2/hTMPRSS2. These discrepancies could potentially result from limitations of the artificial system employed by Meng to directly access virus entry, mostly related to an envelope composition, lacking the main viral membrane protein, and to a different nucleocapsid composition that imposes different interactions to the S transmembrane portion.

Previous studies reported, *in vitro* and *in vivo*, differences in the replication capacity of ancestral SARS-CoV-2 D614G when compared to VOIs and among VOCs^22,24,29,30,32,36,37^. Overall, BA.1 replicates to lower titers in epithelial lung and intestinal cells, primary lung cells, and *ex-vivo* tissue. There are some heterogeneities when comparing ancestral D614G, Alpha, Beta, Gamma, and Delta VOCs and other VOIs replication capacity, but higher replication titers for Alpha, Gamma, and Delta had been reported^29,32,36,37^. Besides evidence suggesting that the attachment/entry steps correlate with these differences, viral factors that could contribute to replicative characteristics of the SARS-CoV-2 variants are lacking. Nchioua^36^ demonstrated that BA.1 is more resistant to Interferons than previous VOCs when replicating in both Calu-3 and iPSC-derived iAT-2 cells. A cell-dependent phenomenon is evident since SARS-CoV-2 replication in bronchial cells *ex-vivo*^29^ and in hamster upper respiratory tract^38^ favors BA.1. Since the physiological temperature of the mammal upper respiratory tract varies between 33-35°C, we reproduced this condition by incubating infected Calu-3 cells at 35°C. The fact that BA.1 was insensitive to temperature change while ancestral D614G, Zeta, Gamma, and Delta had an average 2log10 drop in infectious titers suggests a role for virus-cell interactions to explain BA.1 replicative advantage in the upper respiratory tract^39^. Virus adaptation to replicate at lower temperatures has been demonstrated for Influenza A. For instance, avian Influenza A with a PB2 polymerase mutation could efficiently replicate in mammalian cells when the temperature was reduced^40^. In contrast, a Nucleoprotein mutation at a conserved phosphorylation site of another Influenza A isolate led to a cold-sensitivity phenotype^41^. This phenomenon observed for Influenza could explain that the differences in Zeta, Gamma, Delta, and BA.1 replicative capacity may be related to the viral RNA replication and assembly.

The lower infectious progeny production from BA.1-infected Calu-3 cells could be due to an inefficient accumulation of the sgRNAs, especially at 6-12 hpi., which in turn coincided with the lowest average of dsRNA puncta in BA.1-infected cells and the lowest number of cells positive for dsRNA at 8 hpi. The fittest variants in Calu-3 cells – Gamma, Delta and Zeta – were the ones that had the highest levels of sgRNA from 6 to 12 hpi., the highest average number of dsRNA puncta per cell (Gamma and Zeta), and also the highest percentage of dsRNA positive cells (Zeta and Delta) at 8 hpi.

The double-membrane vesicles (DMVs), which are sites for SARS-CoV-2 viral RNA replication are formed by the association of the viral Nsp3 and Nsp4 with the Endoplasmic Reticulum (ER) and the induction of the ER membrane curvature, generating these RNA replication organelles (ROs). Cryo-tomography analyses showed that Nsp3 and Nsp4 are sufficient to form a pore in the ROs which will serve to translocate the newly synthesized viral genomic RNA and the sgRNAs from the DMV lumen to the cytoplasm^42,43^ to be assembled into new ribonucleocapsid structures, and to be translated, respectively^43^. Cryo-tomography analyses also characterized the inner average diameter of the DMV as 338 nm^42^, which implies an average area of 89.68 μM^2^. Indeed, we observe equivalent puncta area average for all variants, except for Delta which had an average size twice higher than the other variants. The data demonstrated that variants inducing higher DMVs average number and size early on infection accumulate higher levels of sgRNA. Our data shows a positive correlation of higher levels of viral RNA replication with higher levels of SARS-CoV-2 structural N protein in infected cells. Moreover, not only did N protein levels vary between BA.1 and all other variants but also the pronounced accumulation of 25-22kDa smaller bands in Calu-3 cells. The appearance of smaller size proteoforms of SARS-CoV-2 N occurs upon its heterologous expression in various systems^44,45^. and has been related to cleavage by cellular Caspases^46^. However, a detailed work demonstrated that N purified from *E. coli* without contamination with cellular proteases underwent “cleavage” generating smaller size proteins of 29 – 25kDa^47^. These authors suggested an unusual lability of N. Also, two works analyzing the SARS-CoV-2 N protein by native mass spectrometry demonstrated the presence of these smaller proteoforms from purified protein expressed in *E. coli*^48^, and from SARS-CoV-2 infected cells^44^, with most of the proteoforms coinciding in both studies. The full-length N has 419 amino acids and 45.8kDa, Lutomski^48^ demonstrated the existence of 5 proteoforms; 4 with a C-terminal cleavage/breakage (N_1-392_ – 42.9kDa, N_1-273_ – 29.4kDa, N_1-220_ – 23.5kDa, N_1-209_ –22.5kDa), and 1 with a N-terminal cleavage/breakage (N_156-419_ – 28.7kDa) (Supplementary Figure 9). This unexpected lability of N can be explained by the presence of a protein domain rich in the serine-arginine (SR) dipeptide sequence (N-SR-rich domain) which is statistically more likely to be present in proteins with shorter than longer *in vivo* half-lives^47,49^. Thus, the SR-rich domain makes N “unstable”. A high degree of conservation amongst all variants was observed at the proposed cleavage/breakage sites. However, for Delta, the R203M and the G215C substitutions, just preceding the N_220_ and N_273_ cleavage/breakage respectively, close to the N-SR-rich domain, which could explain the almost absence of N proteoforms during Delta infection. Interestingly, except for Delta, Beta and related VOIs, all variants derived from the ancestral D614G fixed the RG203/204KR substitution close to the N-SR-rich domain which has been related to an increase in N phosphorylation when compared to the RG original sequence^50^. When we inspected the N expression in Calu-3 cells infected with a Wuhan-related SARS-CoV-2 A2 isolate, we observe the presence of a small amount of the single proteoform of 23kDa (Supplementary figure 8), suggesting that the KR substitution could contribute to an additional degree of instability of N. Although all the C-terminal proteoforms can bind to the viral genomic RNA a disequilibrium could be detrimental for virus replication, since only dimmers of the full-length protein can bind to the viral genomic RNA for assembly^48^. For instance, N and Nsp3 interaction at the sites of viral RNA replication is necessary for process efficiency^51^. The presence of N could account for ROs maintenance and efficiently sort out the RNA species being translocated from the interior of the DMVs to sites of virus assembly. Thus, the apparent greater instability of the full-length N during the BA.1 replication could explain its reduced ability to replicate in Calu-3 cells. In addition, small amounts of the N proteoforms were observed in Vero-hACE2/hTMPRSS2 cells for all the variants, including BA.1, which replicate efficiently in this cell line. As demonstrated for the Nucleoprotein mutation at a conserved phosphorylation site in Influenza A, which reduced protein oligomerization leading to a cold-sensitivity phenotype^41^, the high sensitivity of B.1.133, Zeta, Gamma and Delta to lower virus replication temperature in Calu-3 could be related to greater N instability due to its supposed less efficient oligomerization, which would not affect more BA.1. BA.1 Nsp3 has unique substitutions in functional relevant domains that could potentially affect its ability to bind N^52^, while the N mutations P13L and ΔERS31-33 residing in the N-arm domain preceding the N-terminal domain (Supplementary Figure 9), which has an RNA binding modulatory role could affect the RNA binding regulation by N. Together it could contribute to a greater N instability.

The determinants of successive selection of SARS-CoV-2 variants are related to a myriad of factors including escaping from immune response elicited by vaccination and/or natural infection, higher transmissibility rates, and intrinsic viral factors. The contribution of intrinsic viral factors is directly related to differences observed in virus replicative capacity among variants and its characterization will contribute to a broader understanding of the epidemic properties of the following emerging SARS-CoV-2 variants. In this sense, this work contributes to demonstrating the importance of the RNA replication step and the role of N in shaping the replicative characteristics of the SARS-CoV-2 variants.

## Material and Methods

### Cell cultures

African green monkey kidney cells-VeroE6, (ATCC CRL-1586) and VeroE6 cells expressing human transmembrane protease serine 2 and human angiotensin-converting enzyme 2-Vero/hACE-2/hTMPRSS2 cells (NR-54970) were maintained in Dulbecco’s Modified Eagle Medium (DMEM), with 4.5 g/L D-glucose (Gibco – cat #11995-073) supplemented with 10% fetal bovine serum (FBS) (Gibco – cat #12657-029), 100 U/ml penicillin, and 100 μg/ml of streptomycin (Gibco – cat #15140-122). Human lung adenocarcinoma epithelial cells - Calu-3, (ATCC HTB-55) were cultured DMEM with 1.0 g/L D-glucose (Gibco – cat #12320-032) supplemented with 10% FBS. All cells were maintained at 37°C and 5% CO_2_. and routinely tested for mycoplasma.

### Viral isolation and viral stocks

SARS-CoV-2 lineages B.1.1.33, Zeta, Gamma, and Delta were isolated from nasopharyngeal swab samples collected from individuals attending the Center for Combating and Studying Emerging and Reemerging Infectious Diseases (NEEDIER) at Federal University of Rio de Janeiro (Rio de Janeiro, Brazil). VeroE6 cells were infected using viral transport medium (VTM) from SARS-CoV-2 positive samples. Briefly, cells were seeded in 6-well plates overnight in DMEM with 10% FBS to achieve 70% confluence. Then, 250 µL of VTM diluted in 250 µL DMEM without FBS was added for one hour. to allow virus adsorption. After incubation, the inoculum was replaced by DMEM with 10% FBS, and cells maintained at 37°C and 5% CO_2_. The culture was observed until visualization of the cytopathic effect (CPE). After this first passage in cell culture, a second passage was performed in the same conditions described above. The second passage in cell culture was titrated and used for viral stock generation. For viral stocks, 2,0 x 10^6^ Vero/hACE-2/hTMPRSS2 (2,0 x 10^6^) cells were plated and in 75 cm^2^ culture flasks and cultured overnight. After incubation, cells were infected with SARS-CoV-2 variants at an (MOI of 0.01) and incubated at 37°C and 5% CO_2_ for 72 h. After this72 h, the supernatant from each infection was harvested and filtered through a 0.22 µM filter. to remove cellular debris. The stocks were aliquoted and stored at -80°C. The Omicron (BA.1) variant was isolated and kindly given by professors Edison Luiz Durigon (USP), Ester Sabino (IMT-SP), Fernando Spilki (FEEVALE-SC), and João Renato Rebello Pinho (HIAE). Omicron BA.1 stock was generated as described above.

### Biosafety

The study was approved by the ethical committee (CAAE-30161620.0.1001.5257) and all the experiments using infectious particles were performed in a BSL-3 laboratory.

### Viral titration

Stocks and experiments supernatants were titrated as follow. Vero/hACE-2/hTMPRSS2 cells were plated overnight in 12-well plates to achieve 90% confluence. Cell media was then replaced by 200 µL of 10-fold serial dilutions of each sample for infection. After one hour of adsorption, media was replaced by DMEM (Gibco – cat # 12100-046) with 1% FBS, 100 u/mL penicillin, 100 µg/mL streptomycin, and 1.4% carboxymethylcellulose (Sigma Aldrich – cat #C5678), followed by incubation for three days at 37 °C with 5% CO_2_. After 3 days of incubation, cells were fixed with 5% formaldehyde and stained with 1% crystal violet in 20% methanol for plaque visualization and quantification. Viral titers were expressed as plaque forming units (PFU) per milliliter.

### Viral binding and entry assay

VeroE6, and Vero/hACE-2/hTMPRSS2 (2,0 x 10^6^ cells), and Calu-3 cells (3,0 x 10^6^ cells) were plated in 24-well plates and infected with SARS-CoV-2 variants with the same viral copy number, diluted in unsupplemented DMEM without FBS, at 4°C for 1h. After viral adsorption, the monolayer was washed with cold 1X PBS to remove unattached particles, and three wells were harvested and submitted to viral RNA extraction with TRIzol (Invitrogen – cat #15596026). Three other wells received DMEM with 10% FBS and were cultured at 37°C with 5% CO_2_. for 1h. After 1h incubation, supernatants and cells were collected for total RNA extraction, using TRIzol for cells and Bio Gene (Bioclin – cat #K204-4) kit for supernatant extraction, respectively, following the manufacturer’s instructions. Detection and quantification of SARS-CoV-2 was made by RT-qPCR using the Detection Kit for 2019 Novel Coronavirus (2019-nCoV) RNA (DaAnGene - cat #DA-930).

### Viral kinetics assay

Viral replication kinetics were performed in 24-well plates seeded with 4,0x10^6^ cells for VeroE6 and Vero/hACE-2/hTMPRSS2, and with 3,5x10^6^ for Calu-3 cells. Monolayers were infected with SARS-CoV-2 variants (MOI of 0.1). Supernatants and cells were collected after 12, 24, 48, and 72 hpi post-infection for Vero cells and up to 96 hpi for Calu-3 cells. The samples were stored at -80. Supernatants were used for viral titration by plaque assay, and RNA extraction with Bio Gene (Bioclin – cat #K204-4). Cells were treated with TRIzol (Invitrogen – cat #15596026)) for RNA extraction.

### cDNA synthesis, genomic and subgenomic detection

For genomic SARS-CoV-2 RNA detection, real-time RT-qPCR reactions were performed using the Detection Kit for 2019 Novel Coronavirus (2019-nCoV) RNA (DaAnGene - cat #DA-930) for detection of N and ORF genes according to manufacturer’s instructions. For subgenomic RNA (sgRNA) and *gapdh* detection, 1 µg of total RNA was used for cDNA synthesis using was performed with the High-Capacity cDNA Reverse Transcription kit (Applied Biosystems – cat # 4368814) priming with random hexamers. N sgRNA was detected using TaqMan™ Universal PCR Master Mix (Applied Biosystems – cat # 4304437) and each reaction contain: 10 µL of TaqMan; 0.08 µL of 100 µM of forward primer (5’ CGA TCT CTT GTA GAT CTG TTC TCT AAA CGA ACT TAT GTA CTC 3’); 0.08 µL of 100 µM of reverse primer (5’ ATA TTG CAG CAG TAC GCA CAC A 3’); 0.04 µL of 100 µM of probe (FAM-5’ACA CTA CTA GCC ATC CTTACT GCG CTT CG 3’-IowaBlack); 6.8 µL of nuclease free water and 3 µL of cDNA. Reactions were cycled as follows: 95°C for 2 minutes; 45 cycles of 95°C for 15 seconds and 53°C for 30 seconds. Sample analysis was performed using a threshold of 0.01.

For *gapdh* was detected using Power SYBR Green PCR master mix reagent (Applied Biosystems – cat # 4309155). Reactions contained: 10 µL of Sybr Green; 0.8 µL of forward primer (5’GTG GAC CTG ACC TGC CGT CT 3’); and 0.8 µL of reverse primer (5’ GGA GGA GTG GGT GTC GTC GT 3’); 7 µL of nuclease free water and 1 µL of cDNA. Reactions were cycled at: 95°C for 2 minutes; 40 cycles of 95°C for 15 seconds and 60°C for 1 minute following the melting cycle. Intracellular sgRNA RNA levels were normalized by *gapdh* cycle threshold values. All reactions were performed at ARIA MX (Agilent) thermocycler.

### SARS-CoV-2 RNA copy number quantification

In order to determine viral RNA copy number a dose-response curve was constructed by simple linear regression. We used a positive control with known gene copies number (2019-nCoV_N_Positive Control, IDT - cat. #10006625) made of an in vitro transcribed and purified plasmid DNA target that contains 200,000 copies of gene N per microliters.

### Cell viability assay

Supernatants from non-infected and infected cell cultures were harvested from all the incubation time points and used for determination of the percentage of cell survival using LDH-Glo™ Cytotoxicity Assay (Promega – cat #J2381) kit following the manufacturer’s instructions.

### Western blot

Whole infected-cell lysates were collected with RIPA buffer (10 mM Tris-Cl [pH 8.0]; 1 mM EDTA; 0.5 mM EGTA; 1% Triton X-100; 0.1% sodium deoxycholate; 0.1% SDS; 140 mM NaCl) added with 0.1% protease inhibitors cocktail (Sigma Aldrich – cat #P8340). Samples were loaded in 4%-20% Mini-Protean gels (Bio-Rad, cat #4561093). The following antibodies were used for viral protein analysis: SARS-CoV-2 anti-Spike and anti-N protein (cat #569996 and #33336, respectively, Cell Signaling) and anti-β-actin (Sigma – cat #a2228); anti-rabbit HRP (Cell Signaling – cat #70745) and anti-mouse HRP (Thermo Scientific – cat #31436).

### Sequencing of viral stocks

All viral stocks were sequenced after cell passage to confirm viral lineage and the emergence of non-defining lineage mutations. The GISAID accession numbers are summarized in Table 1. The complete coverage of the SARS-CoV-2 genome was obtained through NGS, using the Ion AmpliSeq SARS-CoV-2 Research Panel (ThermoFisher Scientific). The multiplex amplification reaction was conducted using 10 µl of the cDNA according to the manufacturer’s instructions for 21 cycles of the multiplex RT-PCR-specific SARS-CoV-2 primers from the panel. Library quantification was performed using the Ion Library TaqMan™ Quantitation Kit (Thermofisher Scientific – cat # 4468802). Libraries were pooled at a concentration of 50pM, and the emulsion PCR and enrichment reactions were conducted on the Ion Chef™ system (ThermoFisher Scientific) using Ion 510, 520 & 530 kits (ThermoFisher – cat # A34461). Using a 530 chip, the sequencing reaction was performed on the Ion S5™ System genetic sequencer (ThermoFisher Scientific). Reads generated were mapped to the SARS-CoV-2 reference genome Wuhan (NCBI GenBank accession number MN908947) using the Ion Browser software included in the Torrent Suite 5.18.1. Virus genome assembly was carried out with the IRMA plugin v.1.3.0.2. Lineage was accessed using the web tool Nextclade version 2.14.1 (Nextstrain project)^27^ through fasta files generated from NGS sequencing.

**Table 1:** Accession number of the SARS-CoV2 sequenced viral stocks at the EpiCoV database from GISAID.

| **SARS-CoV-2 Variant**<br>**and lineage** | **Type Cell and Passage**<br>**Number** | **GISAID number** |
| --- | --- | --- |
| A2 | Vero E6 - 2 | EPI_ISL_528539 |
| B.1.1.33 | original sample (VTM) | EPI_ISL_11836071 |
| B.1.1.33 | Vero/hACE-2/hTMPRSS2- 4 | EPI_ISL_18433588 |
| Zeta P.2 | Vero/hACE-2/hTMPRSS2 - 4 | EPI_ISL_18433589 |
| Gamma P.1 | Vero E6 - 2 | EPI_ISL_18433882 |
| Gamma P.1 | Vero/hACE-2/hTMPRSS2 - 3 | EPI_ISL_18433590 |
| Gamma P.1 | Vero/hACE-2/hTMPRSS2 - 4 | EPI_ISL_18433591 |
| Delta | Vero/hACE-2/hTMPRSS2 - 3 | EPI_ISL_18433883 |
| Delta | Vero/hACE-2/hTMPRSS2 - 4 | EPI_ISL_18433592 |
| Omicron (BA.1) | original sample (VTM) | EPI_ISL_7699344 |
| Omicron (BA.1) | Vero/hACE-2/hTMPRSS2 - 1 | EPI_ISL_18433884 |
| Omicron (BA.1) | Vero/hACE-2/hTMPRSS2 - 2 | EPI_ISL_18433885 |
| Omicron (BA.1) | Vero/hACE-2/hTMPRSS2 - 3 | EPI_ISL_18433886 |
| Omicron (BA.1) | Vero/hACE-2/hTMPRSS2 - 4 | EPI_ISL_18433887 |

### Competition assay

Vero/hACE-2/hTMPRSS2 (4,0 x 10^6^ cells) and Calu-3 cells (3,5 x 10^6^ cells) were seeded in 48-well plates and co-infected with three different proportions of SARS-CoV-2 variants: 1:1, 1:9 and 9:1 resulting in a final MOI of 0.1 PFU. The combination of variants used in this assay was: Gamma and Omicron; Delta and Omicron; Zeta and Gamma; Gamma and Delta. The culture supernatants were harvested after 24 and 48h of infection, and stored at -80 for further RNA extraction. Total RNA was extracted from supernatants with Bio Gene (Bioclin - cat #K204-4) followed by and then used for major and minor viral population detection by NGS or RT-qPCR. For RT-qPCR detection, primers and probes were designed to detect two variant defining mutations: a deletion at NSP6 gene that occurs exclusively at the Gamma variant; and the deletion of aminoacids 69 and 70 at the S gene that occurs in the Alpha and Omicron BA.1 variants. In addition, primers and probes for envelope (E) and nucleocapsid (N) targets were used for total RNA quantification. Samples double negative for NSP6 and 69-70 deletions were classified as Delta or Zeta, while samples positive only for NSP6 deletion were classified as Gamma. The percentage of each variant was calculated as the ΔCt of the specific target minus the nucleocapsid target.

### SARS-CoV-2 whole genome sequencing of the competition assay

For the competition assays revealed by NGS, the SARS-CoV-2 whole genome was recovered using Illumina COVIDSeq Test (Illumina, CA, USA) and the ARTIC 4.1v primer set (https://artic.network/). The final library was submitted to the MiSeq platform (Illumina, CA, USA) to achieve 2000x coverage and the results obtained in the Illumina Basespace Sequence Hub. The reads were assembled using the reference genome MN985325.1 (USA/WA1/2020). S Bioinformatic analysis was performed with Illumina® DRAGEN COVID Lineage App (version 3.5.12) and also assembled by ViralFlow 1.0.0^28^. The ViralFlow pipeline is able to generate a report which includes the analysis of minor variants. For lineage assignment, the Pango Lineage tool was used. Consensus fasta sequences were uploaded to the EpiCoV database in GISAID.

### Immunofluorescence

Calu-3 cells were cultured overnight onto coverslips followed by infection with SARS-CoV-2 variants (at an MOI of 0.1). Then, culture supernatants were harvested at 8- and 24-hours post-infection and cells were fixed with 4% PFA in PBS for 15 min, washed twice with PBS, and permeabilized with 0.1% Triton X-100 (Sigma-Aldrich – X100-5ML) with 3% BSA (Sigma-Aldrich – A1906) in PBS for 20 min. Cells were incubated with J2 anti-dsRNA IgG2a monoclonal antibody (Scicons – cat #RNT-SCI-10010200) diluted 1:1000 in 3% BSA in PBS for 1 h at 4 °C. Then, cells were washed twice in PBS followed by and incubated with AlexaFluor 488-conjugated anti mouse IgG (Thermo Scientific – cat #A28175.) and for diluted 1:1000 in PBS for 40 min and washed three times with PBS followed by incubation with DAPI staining (Invitrogen – cat #D1306) for 10 min. Then, coverslips were mounted with ProLong™ Gold Antifade Reagent (Thermo Scientific – cat #P36934) and imaged on a Zeiss LSM 710 confocal microscope.

### Quantification and statistical analysis

Statistical assessments were conducted for comparing two experimental groups via unpaired multiple *t*-tests. For comparisons involving three or more experimental groups, One-way ANOVA followed by Dunnett’s multiple comparisons or Two-way ANOVA followed by Bonferroni’s multiple comparisons test was employed. Differences were considered statistically significant when p<0.05. Data analysis was executed utilizing GraphPad Prism v 9.4.1.

## Acknowledgments

We thank Mr. Ronaldo Rocha and Mr. Manoel Itamar do Nascimento for technical assistance. We would also like to thank Dr. Andre Felipe Andrade dos Santos, Dr. Atila Duque Rossi, Dr. Mirela D’arc, Dr. Francine Bittencourt Schiffler, Dr Filipe Romero Rebello Moreira, Mr. Marcelo Calado de Paula Tôrres, Mr. Matheus Augusto Calvano Cosentino, Ms. Raíssa Mirella dos Santos Cunha da Costa, and Ms. Thamiris dos Santos Miranda for handling sequencing and sequence analysis.

## Author contributions

I.C, M.R.M.S and G.P.D.S: research design, carried out experiments, analyzed and discussed results. I.C and M.R.M.S manuscript writing. G.P.D.S assemble of figures. A.B.V.M.P western blotting experiments. P.T.C western blotting experiments and discussed results. M.S.C carried out experiments and discussed results. D.M sample collection, handling and storage coordination. R.M.B RT-qPCR for viral competition assay. S.M.C and M.C.C.S NGS sequencing and analysis of viral stocks. V.A.O and R.M.G sample and data collection. E.C.P, M.M.S and P.C.R NGS sequencing and analysis for viral competition assay. D.S.F, R.S and T.M.P.P.C discussed results and manuscript review. AT and LC: research conception and design, study implementation, data analyses and discussion, and manuscript supervision and editing. All authors contributed to the article and approved the submitted version.

## Declaration of interest

The authors declare no conflicts of interest.

## Notes

### Competing Interest Statement

The authors have declared no competing interest.

## References

1 Plante JA, Mitchell B, Plante KS, Debbink K, Weaver SC, Menachery VD. The variant gambit: COVID-19’s next move. Cell Host & Microbe. 2021 Apr 1;29(4):508–15.

2 Van Dorp L, Acman M, Richard D, Shaw L, Ford C, Ormond L, et al. Emergence of genomic diversity and recurrent mutations in SARS-CoV-2. Infection, Genetics and Evolution. 2020 Sep 1;83:104351.

3 Gupta S, Gupta D, Bhatnagar S. Analysis of SARS-CoV-2 genome evolutionary patterns. Microbiology Spectrum. 2024 Jan 10. 10.1128/spectrum.02654-23

4 Korber B, Fischer W, Gnanakaran S, Yoon H, Theiler J, Abfalterer W, et al. Tracking Changes in SARS-CoV-2 Spike: Evidence that D614G Increases Infectivity of the COVID-19 Virus. Cell (Cambridge). 2020 Aug 1;182(4):812–827.e19.

5 Hou YJ, Chiba S, Halfmann P, Ehré C, Kuroda M, Dinnon KH, et al. SARS-CoV-2 D614G variant exhibits efficient replication ex vivo and transmission in vivo. Science (New York, NY). 2020a Dec 18;370(6523):1464–8.

6 Plante JA, Liu Y, Liu J, Xia H, Johnson BA, Lokugamage KG, et al. Spike mutation D614G alters SARS-CoV-2 fitness. Nature. 2020 Oct 26;592(7852):116–21.

7 WHO. Updated working definitions and primary actions for SARSCoV2 variants. Available from: https://www.who.int/publications/m/item/historical-working-definitions-and-primary-actions-for-sars-cov-2-variants.

8 Campbell F, Archer BN, Laurenson-Schafer H, Jinnai Y, Konings F, Batra N, et al. Increased transmissibility and global spread of SARS-CoV-2 variants of concern as at June 2021. Euro Surveillance. 2021 Jun 17;26(24).

9 CoVariants. CoVariants. Available from: https://covariants.org/cases

10 Viana R, Moyo S, Amoako DG, Tegally H, Scheepers C, Althaus CL, et al. Rapid epidemic expansion of the SARS-CoV-2 Omicron variant in southern Africa. Nature (London). 2022 Jan 7;603(7902):679–86.

11 Dhama K, Nainu F, Frediansyah A, Yatoo MI, Mohapatra RK, Chakraborty S, et al. Global emerging Omicron variant of SARS-CoV-2: Impacts, challenges and strategies. Journal of Infection and Public Health. 2023 Jan 1;16(1):4–14.

12 Da Silva Cândido D, Claro IM, De Jesus JG, De Souza WM, Moreira FRR, Dellicour S, et al. Evolution and epidemic spread of SARS-CoV-2 in Brazil. Science. 2020 Sep 4;369(6508):1255–60.

13 Naveca FG, Nascimento VAD, De Souza VC, De Lima Guerra Corado A, Nascimento F, Silva G, et al. COVID-19 in Amazonas, Brazil, was driven by the persistence of endemic lineages and P.1 emergence. Nature Medicine. 2021 May 25;27(7):1230–8.

14 Faria NR., Claro IM., Candido DD., Franco LA., Andrade PS., Coletti, TM., et al. Genomic characterization of an emergent SARS-CoV-2 lineage in Manaus: preliminary findings. 2021. Artic Network.

15 Naveca FG., Nascimento V., Souza V., Corado A., Nascimento F., Silva G., et al. Phylogenetic relationship of SARS-CoV-2 sequences from Amazonas with emerging Brazilian variants harboring mutations E484K and N501Y in the Spike protein. 2021. Artic Network.

16 Voloch CM, Da Silva Francisco R, De Almeida LGP, Cardoso CC, Brustolini OJB, Gerber AL, et al. Genomic Characterization of a Novel SARS-CoV-2 Lineage from Rio de Janeiro, Brazil. Journal of Virology. 2021 Apr 26;95(10).

17 Dashboard-en Genomahcov - Fiocruz. Avilable from: https://www.genomahcov.fiocruz.br/dashboard-en/.

18 Li B, Deng A, Li K, Hu Y, Li Z, Shi Y, et al. Viral infection and transmission in a large, well-traced outbreak caused by the SARS-CoV-2 Delta variant. Nature Communications. 2022 Jan 24;13(1).

19 Moreira FRR, D’arc M, Mariani D, Herlinger AL, Schiffler FB, Rossi ÁD, et al. Epidemiological dynamics of SARS-CoV-2 VOC Gamma in Rio de Janeiro, Brazil. Virus Evolution. 2021 Oct 1;7(2).

20 Ong SWX, Chiew CJ, Ang LW, Mak TM, Cui L, Toh MPHS, et al. Clinical and virological features of SARS-COV-2 variants of concern: A retrospective cohort study comparing B.1.1.7 (Alpha), B.1.315 (Beta), and B.1.617.2 (Delta). Social Science Research Network. 2021 Jan 1.

21 Shuai H, Chan JF, Hu B, Chai Y, Yuen TT, Yin F, et al. Attenuated replication and pathogenicity of SARS-CoV-2 B.1.1.529 Omicron. Nature (London). 2022 Jan 21;603(7902):693–9.

22 Hui K, Ho JW, Cheung MC, Ng K, Ching RHH, Lai KL, et al. SARS-CoV-2 Omicron variant replication in human bronchus and lung ex vivo. Nature (London). 2022a Feb 1;603(7902):715–20.

23 Halfmann P, Iida S, Iwatsuki-Horimoto K, Maemura T, Kiso M, Scheaffer SM, et al. SARS-CoV-2 Omicron virus causes attenuated disease in mice and hamsters. Nature (London). 2022 Jan 21;603(7902):687–92.

24 Peacock TP, Brown JC, Zhou J, Thakur N, Newman J, Kugathasan R, et al. The altered entry pathway and antigenic distance of the SARS-CoV-2 Omicron variant map to separate domains of spike protein. bioRxiv (Cold Spring Harbor Laboratory). 2022 Jan 3.

25 Laitman AM, Lieberman JA, Hoffman NG, Roychoudhury P, Mathias PC, Greninger AL. The SARS-COV-2 Omicron variant does not have higher nasal viral loads compared to the Delta variant in symptomatic and asymptomatic individuals. Journal of Clinical Microbiology. 2022 Apr 20;60(4).

26 Sentis C, Billaud G, Bal A, Frobert É, Bouscambert M, Destras G, et al. SARS-CoV-2 Omicron Variant, Lineage BA.1, Is Associated with Lower Viral Load in Nasopharyngeal Samples Compared to Delta Variant. Viruses. 2022 Apr 28;14(5):919.

27 Aksamentov I, Roemer C, Hodcroft EB, Neher RA. Nextclade: clade assignment, mutation calling and quality control for viral genomes. Journal of Open Source Software. 2021 Nov 30;6(67):3773.

28 Dezordi FZ, Da Silva Neto AM, De Lima Campos T, Jerônimo PMC, Aksenen CF, De Almeida SC, et al. ViralFlow: a versatile automated workflow for SARS-COV-2 genome assembly, lineage assignment, mutations and intrahost variant detection. Viruses. 2022 Jan 23;14(2):217.

29 Hui K, Ng K, Ho JCW, Yeung HW, Ching RHH, Gu H, et al. Replication of SARS-CoV-2 Omicron BA.2 variant in ex vivo cultures of the human upper and lower respiratory tract. EBioMedicine. 2022b Sep 1;83:104232.

30 Begum MM, Ichihara K, Takahashi O, Nasser H, Jonathan M, Tokunaga K, et al. Virological characteristics correlating with SARS-CoV-2 spike protein fusogenicity. Frontiers in Virology. 2024 Mar 14;4.

31 Willett BJ, Grove J, MacLean OA, Wilkie C, De Lorenzo G, Furnon W, et al. SARS-CoV-2 Omicron is an immune escape variant with an altered cell entry pathway. Nature Microbiology. 2022 Jul 7;7(8):1161–79.

32 Meng B, Abdullahi A, Ferreira I, Goonawardane N, Saito A, Kimura I, et al. Altered TMPRSS2 usage by SARS-CoV-2 Omicron impacts infectivity and fusogenicity. Nature. 2022 Feb 1;603(7902):706–14.

33 Melikyan GB, Markosyan RM, Hemmati H, Delmedico MK, Lambert DM, Cohen FS. Evidence That the Transition of HIV-1 Gp41 into a Six-Helix Bundle, Not the Bundle Configuration, Induces Membrane Fusion. The Journal of Cell Biology. 2000 Oct 16;151(2):413–24.

34 Henderson HI., & Hope TJ. The temperature arrested intermediate of virus-cell fusion is a functional step in HIV infection. 2006. Virology journal, 3, 36.

35 Chou T. Stochastic Entry of Enveloped Viruses: Fusion versus Endocytosis. Biophysical Journal. 2007 Aug 1;93(4):1116–23.

36 Nchioua R, Schundner A, Klute S, Koepke L, Hirschenberger M, Noettger S, et al. Reduced replication but increased interferon resistance of SARS-CoV-2 Omicron BA.1. Life Science Alliance. 2023 Mar 28;6(6):e202201745.

37 Mautner L, Hoyos M, Dangel A, Berger C, Ehrhardt A, Baiker A. Replication kinetics and infectivity of SARS-CoV-2 variants of concern in common cell culture models. Virology Journal. 2022 Apr 26;19(1).

38 Escalera A, González-Reiche AS, Aslam S, Mena I, Laporte M, Pearl R, et al. Mutations in SARS-CoV-2 variants of concern link to increased spike cleavage and virus transmission. Cell Host & Microbe. 2022 Mar 1;30(3):373–387.e7.

39 Stauft CB, Sangare K, Wang TT. Differences in new variant of concern replication at physiological temperatures in vitro. The Journal of Infectious Diseases (Online University of Chicago Press. 2022 Jun 27;227(2):202–5.

40 Hayashi T, Wills S, Bussey KA, Takimoto T. Identification of influenza A virus PB2 residues involved in enhanced polymerase activity and virus growth in mammalian cells at low temperatures. Journal of Virology. 2015 Aug 1;89(15):8042–9.

41 Zheng W, Cui L, Li M, Li Y, Fan W, Yang L, et al. Nucleoprotein phosphorylation site (Y385) mutation confers temperature sensitivity to influenza A virus due to impaired nucleoprotein oligomerization at a lower temperature. Science China Life Sciences. 2020 Aug 11;64(4):633–43.

42 Klein S, Cortese M, Winter SL, Wachsmuth-Melm M, Neufeldt CJ, Cerikan B, et al. SARS-CoV-2 structure and replication characterized by in situ cryo-electron tomography. Nature Communications. 2020 Nov 18;11(1).

43 Zimmermann L, Zhao X, Makroczyová J, Wachsmuth-Melm M, Bartenschlager R, Hensel Z, et al. SARS-CoV-2 nsp3 and nsp4 are minimal constituents of a pore spanning replication organelle. Nature Communications. 2023 Nov 30;14(1).

44 Meyer B, Chiaravalli J, Gellenoncourt S, Brownridge P, Bryne DP, Daly LA, et al. Characterising proteolysis during SARS-CoV-2 infection identifies viral cleavage sites and cellular targets with therapeutic potential. Nature Communications. 2021 Sep 21;12(1).

45 Emmott E, Munday DC, Bickerton E, Britton P, Rodgers M, Whitehouse A, et al. The cellular interactome of the coronavirus infectious bronchitis virus nucleocapsid protein and functional implications for virus biology. Journal of Virology. 2013 Sep 1;87(17):9486–500.

46 Diemer C, Schneider M, Seebach J, Quaas J, Frösner G, Schätzl H, et al. Cell Type-Specific Cleavage of Nucleocapsid Protein by Effector Caspases during SARS Coronavirus Infection. Journal of Molecular Biology/Journal of Molecular Biology. 2008 Feb 1;376(1):23–34.

47 Mark JK, Li X, Cyr TD, Fournier S, Jaentschke B, Hefford MA. SARS coronavirus: Unusual lability of the nucleocapsid protein. Biochemical and Biophysical Research Communications. 2008 Dec 1;377(2):429–33.

48 Lutomski CA, El-Baba TJ, Bolla JR, Robinson CV. Multiple Roles of SARS-CoV-2 N Protein Facilitated by Proteoform-Specific Interactions with RNA, Host Proteins, and Convalescent Antibodies. JACS Au. 2021 Jun 15;1(8):1147–57.

49 Guruprasad K, Reddy BVB, Pandit MW. Correlation between stability of a protein and its dipeptide composition: a novel approach for predicting in vivo stability of a protein from its primary sequence. Protein Engineering, Design & Selection. 1990 Jan 1;4(2):155–61.

50 Johnson BA, Zhou Y, Lokugamage KG, Vu MN, Bopp NE, Crocquet-Valdes PA, et al. Nucleocapsid mutations in SARS-CoV-2 augment replication and pathogenesis. PLOS Pathogens. 2022 Jun 21;18(6):e1010627.

51 Li P, Xue B, Schnicker N, Wong LYR, Meyerholz DK, Perlman S. Nsp3-N interactions are critical for SARS-CoV-2 fitness and virulence. Proceedings of the National Academy of Sciences of the United States of America. 2023a Jul 24;120(31).

52 Hossain A, Trishna SA, Rashid AA, Khair S, Alam ASMRU. Unique mutations in SARS-CoV-2 Omicron subvariants’ non-spike proteins: Potential impacts on viral pathogenesis and host immune evasion. Microbial Pathogenesis. 2022 Sep 1;170:105699.

